# Antiviral activity of Cytomegalovirus-specific CD4+ T cells against lytically infected non-haemopoietic cells

**DOI:** 10.1101/2022.12.08.519485

**Authors:** EY Lim, CJ Houldcroft, SE Jackson, G Okecha, M Wills

## Abstract

HCMV establishes a lifelong latent infection in its host, eliciting a broad immune response involving both the innate and adaptive arms of immunity. In immunocompromised individuals, recovery of the HCMV-specific CD4+ T cell response is associated with a lower risk of HCMV reactivation and disease, suggesting that CD4+ T cells contribute significantly to control of viral replication *in vivo*. However, most prior studies of the HCMV-specific CD4+ T cell response have been performed with the aim of either predicting those at risk for HCMV reactivation and end-organ disease, or identifying markers which are associated with higher risk of HCMV viraemia. Few studies have examined the mechanisms of viral control of CD4+ T cells in response to cells lytically infected with HCMV. In this paper, we show that CD4+ T cells from a HCMV-seropositive donor can prevent viral dissemination in autologous fibroblasts, that the secretome from these cells is also antiviral, and that IFN-γ is a key cytokine in the secretome. We also show that MHC Class II antigen presentation by monocytes was critical to driving this HCMV-specific CD4+ T cell response.

## INTRODUCTION

In immunocompetent individuals, Human cytomegalovirus (HCMV) infection usually manifests as a largely self-resolving viral illness, as the innate and adaptive arms of the immune system limit viral replication and prevent end-organ disease.^1^ The virus induces a robust cell-mediated response, including both CD4+ and CD8+ T cells. In HCMV-seropositive individuals, CD8+ T cells have been found to respond to up to 213 open reading frames (ORF), while CD4+ T cells have been found to recognise up to 125 different ORFs, although there is a wide variability in the size of these T cell responses between individuals.^2^ As with all herpesviruses, HCMV has two life cycle phases: a lytic phase with the production of new virions and a latent phase where there is a restriction gene transcription profile and no new virion production. Following primary infection, HCMV establishes latency predominantly in the CD34+ haematopoietic stem cell population in the bone marrow^3,4^ and into the myeloid lineage CD14+ monocytes released into the peripheral blood and can thus migrate into tissues.^5–7^

HCMV encodes multiple immune evasion proteins that allows the virus to modulate intrinsic, innate and adaptive immune responses, the end result of this being the persistence of active primary infection viraemia, which is accompanied by virus excretion for months (in adults) or even years (in children).^8^ So far, more than 40 HCMV gene products have been identified to have a role in modulating the immune response in lytically infected cells.^9–11^ HCMV is able to periodically reactivate, even in the presence of memory immune response (likely due to immune evasion genes) leading to a lytic infection. In a healthy host, periods of reactivation and viral shedding occur, but are ultimately controlled by the immune response.^12^ However, in an immunocompromised host, the lack of adequate anti-viral effector function by a compromised immune system can result in persistent viral replication and eventual end-organ disease. As a result, HCMV is one of the most common complications affecting both solid-organ and haematopoietic stem cell transplants in the paediatric and adult populations.^13,14^ It is particularly notable that persistent viral shedding into the urine and saliva of HCMV-infected but otherwise healthy children is associated with a lack of a HCMV-specific CD4+ T cell response,^15^ and that episodes of recurrent viraemia and end-organ disease among transplant recipients are also associated with a poorer HCMV-specific CD4+ T cell response,^16–19^ suggesting that CD4+ T cells contribute to the prevention of unchecked viral replication and end-organ disease in an immunocompetent host. Multiple studies in both solid organ and haematopoietic stem cell transplant recipients have shown that the recovery of CD4+ T cell response in these recipients was associated with a lower risk of HCMV reactivation and disease.^20^

In haematopoietic stem cell and solid organ transplant recipients, examination of the CD4+ T cell response to HCMV has largely been performed with the aim of either predicting those at risk for HCMV reactivation and end-organ disease, or identifying markers which are associated with higher risk of HCMV viraemia. A few studies have also examined the phenotypic markers and cytokine production by these HCMV-specific CD4+ T cells, with evidence of higher occurrences of CD107a, CD40L, and loss of CD27 and CD28 in these cells.^21–26^ The majority of these studies have relied on the production of interferon-γ (IFN-γ) by CD4+ T cells in response to peptide stimulation or virally-infected dendritic cells as a marker for predicting the effectiveness of HCMV-specific CD4+ T cell responses. However, this does not make an assessment of the antiviral capacity of CD4+ T cells in response to cells lytically infected with HCMV.

Previously, we showed that CD4+ T cells from a HCMV seropositive donor co-cultured with autologous HCMV-infected dendritic cells and fibroblasts prevented CMV dissemination in the fibroblasts.^27^ In this manuscript, we sought to investigate if CD4+ T cells from healthy HCMV-seropositive donors were able to control HCMV replication in HCMV-infected autologous fibroblasts, a cell line that does not constituently express MHC Class II. We demonstrate that CD4+ cells from HCMV-seropositive donors were able to limit immediate-early (IE) and late CMV gene expression in vitro and that the secretome from these CD4+ T cells was also anti-viral when transferred onto a fresh culture of CMV infected cells. Analysis of this secretome demonstrated multiple antiviral cytokines and chemokines important for recruitment of T cells, NK cells and myeloid cells. In particular, there was a substantial IFN-γ response. Given that fibroblasts do not constitutively express MHC Class II, it was unclear how presentation of HCMV antigens to CD4+ T cells was occurring. We subsequently found that critical to driving the HCMV specific response was MHC Class II antigen presentation by monocytes leading to IFN-γ expression and subsequent induction of MHC Class II expression by fibroblasts allowing these cells to be directly recognized by CD4+ T cells. We also found that IFN-γ present in these anti-viral secretomes was able to induce MHC Class II presentation on endothelial and epithelial cells, which are additional important cell types supporting HCMV replication during HCMV end-organ disease.^28^

## RESULTS

### CD4+ cells from HCMV-seropositive but not HCMV-seronegative donors are able to control viral dissemination in an autologous in vitro assay

Previous publications from our group have demonstrated the use of a viral dissemination assay (VDA) to quantify the amount of antiviral activity mediated by either whole PBMCs, CD8+ T cells or NK cells in an autologous in vitro assay, using a clinical isolate of HCMV strain Merlin which expresses mCherry during the immediate-early (IE) phase of lytic infection and GFP at the late phase of lytic infection^29–32^ (Supp Fig.1). We therefore sought to investigate if CD4+ cells from HCMV-seropositive donors were able to exert direct antiviral activity on a cell type that supports full lytic HCMV infection but does not constitutively express MHC Class II. CD4+ cells from 13 healthy HCMV seropositive and 10 healthy seronegative donors were isolated from PBMCs using anti-CD4 microbeads, followed by magnetic column separation. These CD4+ cells were then added to autologous fibroblasts that had been infected with HCMV at a low multiplicity of infection (MOI) and co-cultured for 7—10 days to allow viral replication and dissemination. Fibroblasts were harvested and analysed by flow cytometry for mCherry and GFP expression to determine the effect of CD4 cells on viral replication and dissemination.

The results show that CD4+ cells from HCMV-seropositive donors were able to control both viral spread (measured by IE gene UL36/mCherry expression) and virus replication (measured by late gene UL32/GFP expression) in an E:T dependent fashion. CD4+ T cell control was dependent on HCMV serostatus as there was a statistically significant difference between the amount of control seen in HCMV-seropositive compared to HCMV-seronegative donors (Fig. 1A). This would suggest that the ability of CD4+ cells to control HCMV infection is related to previous infection with HCMV and consistent with the presence of antigen-specific memory CD4+ T cells. To demonstrate the presence of HCMV-specific CD4+ T cell memory responses, PBMC depleted of CD8+ T cells from the same HCMV-seropositive and HCMV-seronegative donors were stimulated with overlapping peptide pools specific to immunodominant HCMV proteins in an IFN-γ FlurosSpot assay and the sum of HCMV-specific responses determined. These peptide pools contained HCMV peptides expressed in both lytic and latent infection (UL138, Latency Unique Nuclear Antigen, US28, vIL-10, UL83, UL144, UL122, UL123, UL82, US3) and were chosen as they had been shown to stimulate the highest IFN-γ responses in a cohort of healthy donors in an earlier publication by our group.^33^ As expected, HCMV seropositive donors showed a higher frequency of HCMV-specific recall CD4+ T cells as compared to naïve HCMV-seronegative donors (Fig 1B).

**Figure 1.**
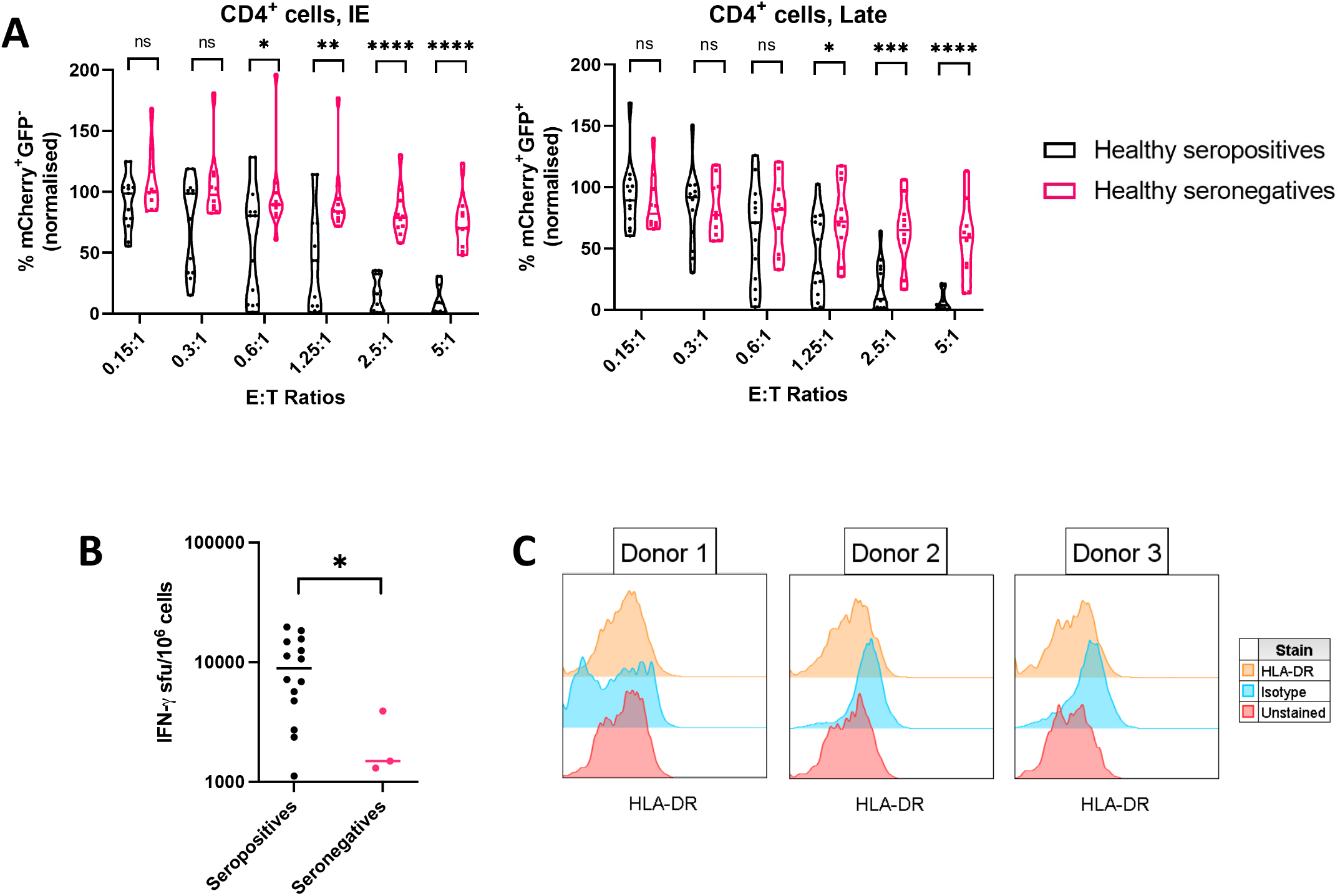
CD4+ cells from HCMV-seropositive donors are able to control early and late viral spread. **(A)** CD4+ cells from 13 healthy HCMV-seropositive and 10 healthy HCMV-seronegative donors were added at starting E:T ratios of 5:1 followed by serial halving dilutions to autologous primary dermal fibroblasts infected with HCMV at 1 dpi and incubated for 7-10 days before harvest and analysis by flow cytometry for amount of cells with IE CMV gene expression (mCherry+GFP-) and late CMV gene expression (mCherry+GFP+). Each data point on the violin plot represents triplicate of wells from one donor. **(B)** PBMCs from 14 healthy HCMV-seropositive and 3 HCMV-seronegative donors were depleted of CD8+ T cells before co-culturing with CMV peptide mixes and incubating for 48 hours before analysis by Fluorospot assay for amount of IFN-γ production. Median of both groups shown. **(C)** Dermal fibroblasts from 3 healthy donors (2 HCMV-seropositive, 1 HCMV-seronegative) were stained for expression of HLA-DR on the cell surface. Statistics performed using Mann Whitney-U test. *p < 0.05, **p < 0.01, ***p < 0.001, ****p < 0.0001.

However, CD4+ T cells recognise antigen via the MHC Class II antigen presentation pathway and MHC Class II is not expected to be constitutively expressed on primary dermal fibroblasts. We confirmed this by staining for MHC Class II HLA-DR on the surface of primary dermal fibroblasts derived from 3 of the donors used in this study (Fig. 1C).

### Secretome from CD4+ cells co-cultured with HCMV-infected autologous fibroblasts is virustatic and contains IFN-γ that can upregulate MHC Class II expression on primary fibroblasts cells

CD4+ T cells can exert their antiviral effector functions by the production of antiviral cytokines as well as via a subset of CD4 cells possessing direct cytotoxic activity.^20^ We next sought to examine the cytokines and chemokines secreted by CD4+ cells following their coculture with HCMV infected autologous fibroblasts, also known as the secretome. The secretome was generated by infecting primary dermal fibroblasts from an HCMV seropositive donor with HCMV, followed by co-incubation with autologous CD4+ cells (Fig.2A). As a control, CD4+ cells from the same donor were added to uninfected fibroblasts. We first determined if this secretome had antiviral activity by transferring it into a HCMV VDA and determining the extent of virus-infected cells 7 days later, by measuring mCherry and GFP expression by flow cytometry. The results very clearly demonstrated that the secretome derived from co-incubation of CD4+ cells with HCMV-infected fibroblasts had anti-HCMV activity which was concentration dependent and not present in the secretome generated from CD4+ cells co-incubated with uninfected cells (Fig 2B). We next examined if this antiviral effect was virustatic or virucidal.

**Figure 2.**
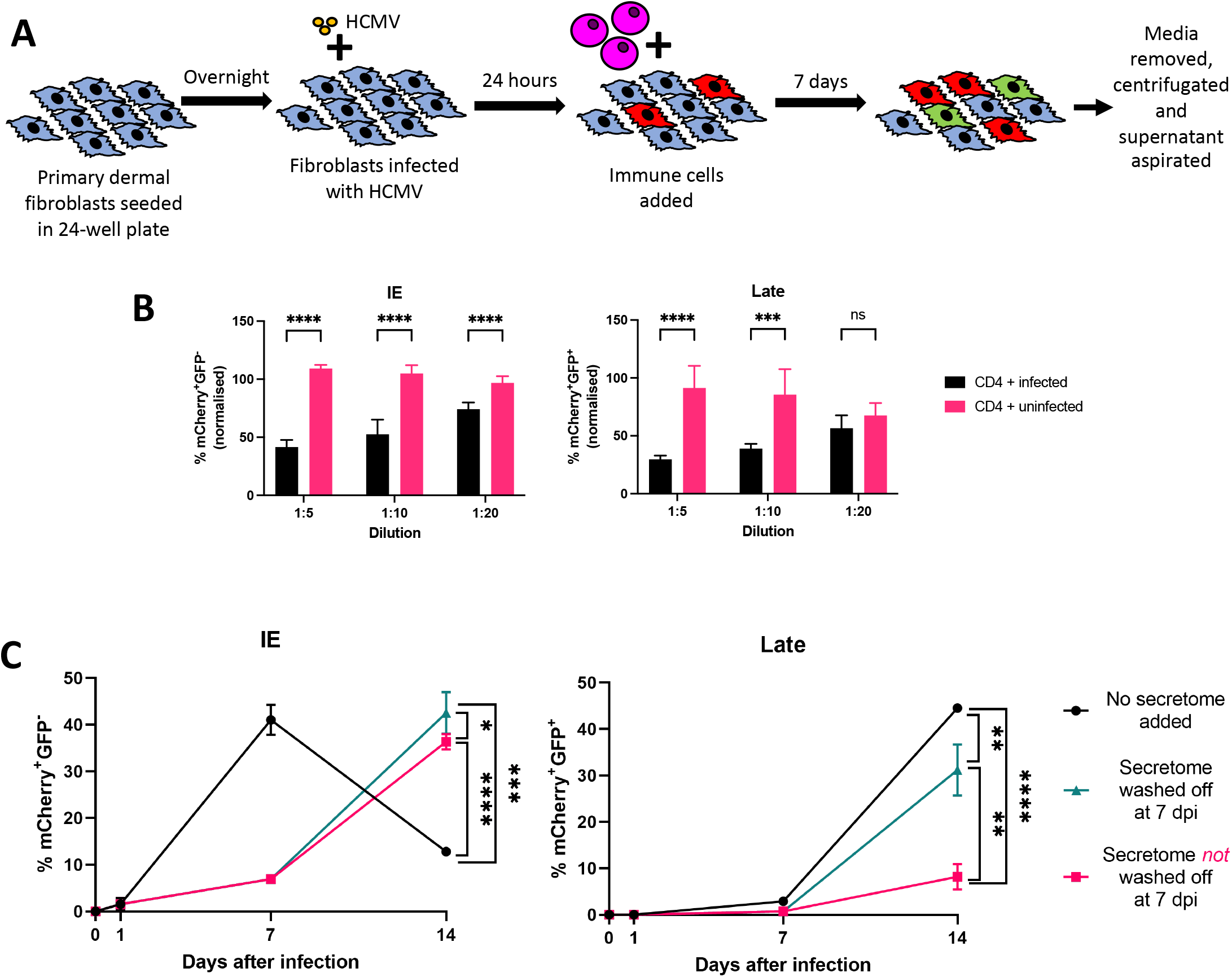
CD4+ cells exert virustatic effects on HCMV-infected fibroblasts via their secretome. **(A)** Secretomes of CD4+ cells or PBMCs from HCMV-seropositive donors co-cultured with HCMV-infected fibroblasts were made by incubating immune cells with HCMV-infected autologous fibroblasts for 7 days before aspiration of media, as illustrated. **(B)** Secretomes from CD4+ cells co-cultured with autologous fibroblasts infected with HCMV (“CD4 + infected”) or uninfected fibroblasts (“CD4 + uninfected”) were added to HCMV-infected fibroblasts (MOI = 0.09) at 1 dpi and incubated for a further 7 days before harvest and analysis by flow cytometry for percentage of cells in IE and late CMV gene expression. **(C)** Secretomes from CD4+ cells co-cultured with HCMV-infected autologous fibroblasts were added to HCMV-infected fibroblasts at 1 day post-infection. These co-cultures were incubated for a further 6 days, whereupon half the wells had secretomes replaced with media (“secretome washed off”) and half did not (“secretome not washed off”). Plates were incubated for a further 7 days. Flow cytometry analysis at days 1, 7 and 14 post-infection show the amount of cells with IE (left panel) or late (right panel) CMV gene expression. Error bars represent SD of triplicates, statistics performed using Student’s *t*-test. *p < 0.05, **p < 0.01, ***p < 0.001, ****p < 0.0001.

Secretomes from CD4+ cells co-cultured with HCMV-infected fibroblasts or media as a control were added to HCMV-infected fibroblasts. Wells were analysed for virus infected cells at 7 dpi by flow cytometry. The wells which had secretome added to them showed an inhibition of both immediate-early and late viral gene expression at 7 dpi (Fig.2C). The secretome was aspirated from duplicate wells and replaced with fresh media, while the secretome was left on a further set of duplicate wells. The assay was then incubated for an additional 7 days before harvest and analysis by flow cytometry. The results show that by 14 dpi, the percentage of mCherry expressing cells was similar between wells with secretome left on or washed off. However, virus late gene expression was still partially inhibited in the wells still containing secretome, while wells that had the secretome removed had significantly higher levels of late gene-expressing cells (Fig.2C). These findings suggest that the secretome is virustatic, but not virucidal, as addition of secretome initially inhibits immediate-early and late CMV gene expression but removal of the secretome led to a resumption of progression of viral infection.

We subsequently proceeded to identify the differentially induced cytokines and chemokines present in secretomes from CD4+ cells co-incubated with either HCMV infected or uninfected autologous fibroblasts. Using a proteomic cytokine array that could detect 105 human cytokines, we showed that CD4+ cells co-incubated with HCMV-infected autologous fibroblasts had upregulated levels of chemokines induced by IFN-γ, such as CXCL11 and CXCL9. Other highly upregulated cytokines included, colony stimulating factors (G-CSF, GM-CSF, IL-3, CSF1), and members of the CC chemokine family (CCL3/CCL4, CCL5, CCL7, CCL20). In addition, IFN-γ and tumour necrosis factor-α (TNF-α) were also upregulated. Other cytokines that were highly upregulated include cytokines involved with macrophage regulation, such as macrophage migration inhibitory factor (MIF), an important regulator of innate immunity,^34^ and macrophage inflammatory protein (MIP-3β), which is produced by dendritic cells^35^ (Fig.3A). Gene ontology analysis of the upregulated cytokines showed that they were most strongly associated with regulation of multicellular processes, in particular regulation of immune system processes and leukocyte migration (Supp Fig.2).

**Figure 3.**
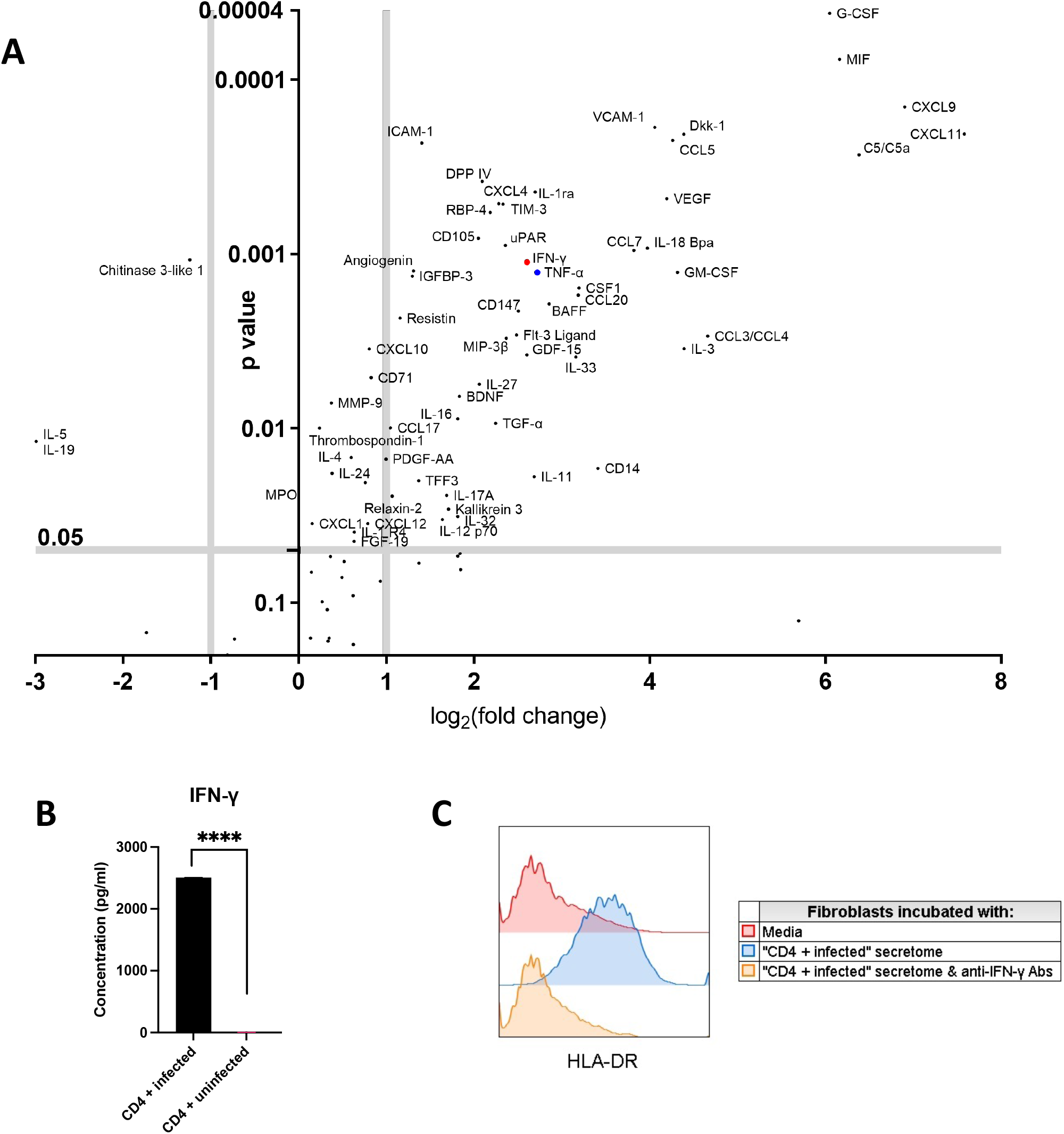
Secretomes of CD4+ cells co-cultured with HCMV-infected fibroblasts contain antiviral cytokines. Secretomes of CD4+ cells co-cultured with either HCMV-infected fibroblasts or uninfected fibroblasts were assayed with a chemiluminescent cytokine array and the relative amounts of various cytokines were compared between the two secretomes. Fold change values represent the amount of cytokines in secretomes of CD4+ cells co-cultured with HCMV-infected fibroblasts versus secretomes of CD4+ cells co-cultured with uninfected fibroblasts. **(B)** LegendPlex analysis of cytokines in “CD4 + infected”, “CD4 + uninfected” secretomes was performed and IFN-γ was the cytokine found to be in the highest concentration. **(C)** Fibroblasts were incubated with secretomes from CD4+ cells co-cultured with HCMV-infected autologous fibroblasts (“CD4 + infected” secretome) in the presence or absence of anti-IFN-γantibodies for 72 hours before harvest and analysis for HLA-DR expression by flow cytometry. Error bars represent SD of duplicate wells, statistics performed using Student’s *t*-test. *p < 0.05, **p < 0.01, ***p < 0.001, ****p < 0.0001

In order to validate a number of the differentially expressed cytokines and chemokines, we also analysed the secretomes using a multiplex bead-based assay to quantify IFN-γ, IFN-α2, IFN-β, IFN-λ1, IFN-λ2/3, TNF-α, interleukin (IL)-1β, IL-8, IL-10 and GM-CSF (Fig. 3B, Supp Fig.3). The results verified that there was induction of IFN-γ, TNF-α and GM-CSF in addition to the immunosuppressive cytokine IL-10. IFN-γ is a type II interferon that regulates a wide range of immune responses and is a key regulator of the T_h_1 response. Importantly, as alluded to in the introduction, it has also been shown to upregulate MHC Class I and II expression and interferon-inducible antiviral mechanisms.^36,37^ Given that IFN-γ was the cytokine found in the highest concentration in the secretome from CD4+ cells co-cultures with HCMV-infected fibroblasts, we thus asked if this secretome could induce MHC Class II expression on the surface of uninfected fibroblasts. This secretome was added to uninfected fibroblasts and incubated for 72 hours and stained for MHC Class II HLA-DR expression, with secretomes from CD4+ cells cocultured with uninfected autologous fibroblasts as a control. The results show a substantial induction of HLA-DR. However, in the presence of neutralizing anti-IFN-γ antibodies, HLA-DR was not induced compared to isotype control (Fig 3C). Therefore, these results demonstrate that IFN-γ present in the secretome is both necessary and sufficient for MHC Class II induction.

### Addition of CD4+ cells to autologous HCMV-infected fibroblasts induces MHC Class II, which is abrogated by blocking IFN-γ

CD4+ T cells recognise antigen via the MHC Class II presentation pathway.^38^ However, as demonstrated earlier, the primary dermal fibroblasts used in our assays do not constituently express MHC Class II. The observation that CD4+ cells could control HCMV gene expression in infected fibroblasts was thus unexpected. Nevertheless, in the results presented so far we have shown that the secretome of CD4+ cells co-cultured with infected autologous fibroblasts induces IFN-γ and this can then stimulate fibroblasts to express MHC Class II. We therefore sought to further investigate whether adding CD4+ cells to HCMV-infected fibroblasts in a low multiplicity of infection (MOI) VDA assay could induce expression of MHC Class II on the cell surface of infected and uninfected fibroblasts, and whether these effects were mediated by IFN-γ.

CD4+ cells from a HCMV-seropositive donor were added to HCMV-infected fibroblasts at an E:T ratio of 1.5:1 at 1 dpi either with or without neutralizing anti-IFN-γ antibodies, and incubated for 7 days before harvest, staining for cell surface HLA-DR expression and analysis by flow cytometry. The results demonstrate that at 7 days of co-culture uninfected fibroblasts express high levels of HLA-DR. In addition, as the fibroblasts become infected and express immediate-early CMV genes (mCherry+) and then late gene expression (GFP+), there was progressively lower cell surface expression of MHC Class II (Fig.4A). The addition of neutralizing anti-IFN-γ antibodies to the cocultures prevented the expression of HLA-DR on the uninfected fibroblasts (Fig.4B).

**Figure 4.**
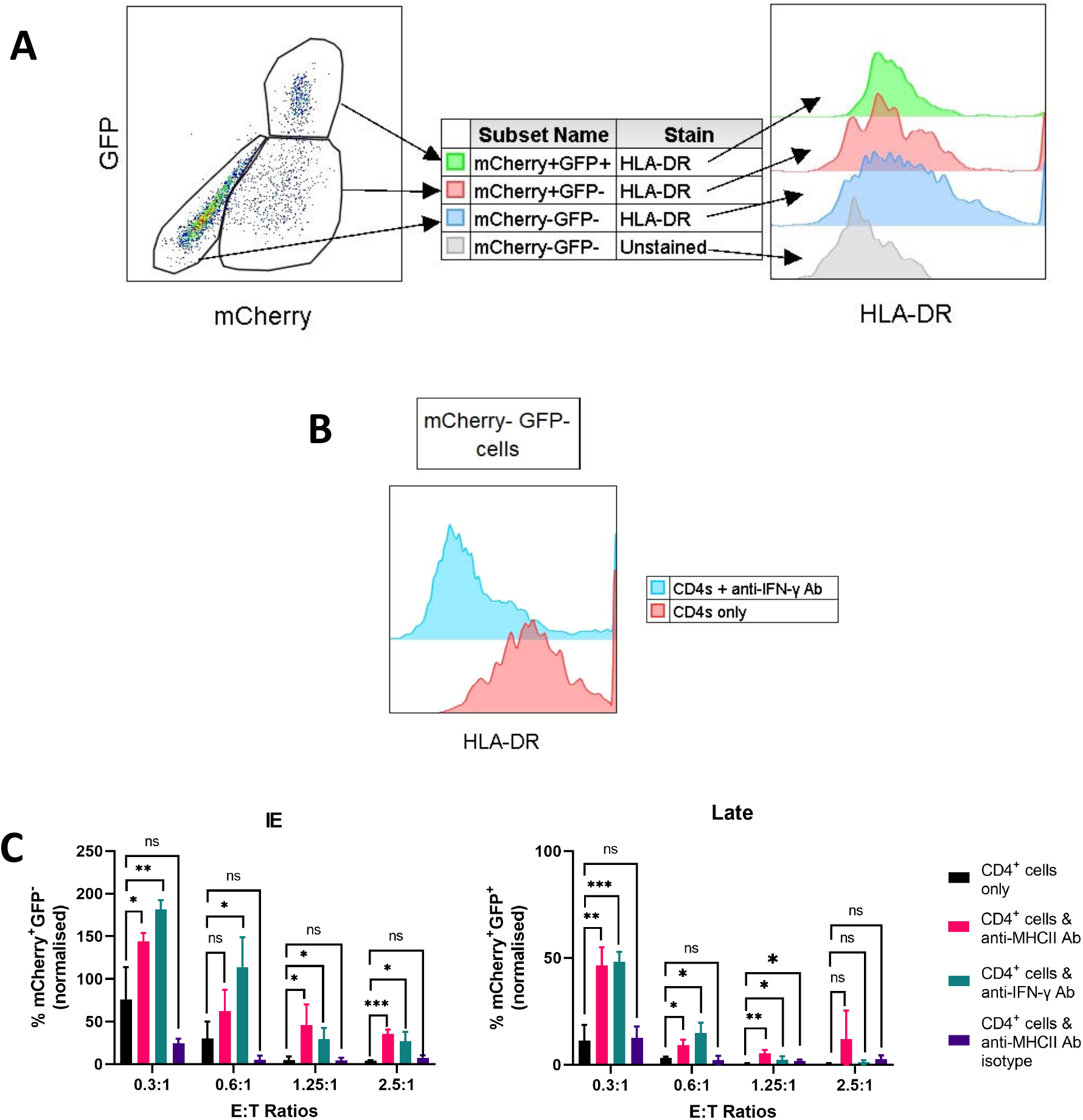
CD4+ cells exert an antiviral effect on HCMV-infected fibroblasts via an MHC Class II pathway, which is mediated by IFN-γ. Primary dermal fibroblasts were infected with HCMV (MOI = 0.04) and CD4+ cells were added at E:T ratio of 1.5:1, with or without anti-IFN-γ antibody, at 1 dpi and incubated for a further 7 days before harvest, staining for HLA-DR and analysis by flow cytometry. **(A)** shows the amount of HLA-DR on the uninfected (mCherry-GFP-), immediate-early CMV gene-expressing (mCherry+GFP-) and late CMV gene-expressing (mCherry+GFP+) populations, with mCherry-GFP-population from an unstained well as a control. **(B)** Histogram plots of the amount of HLA-DR expression on the mCherry-GFP-populations from a well which had CD4+ cells only, and a well which had CD4+ cells and anti-IFN-γ antibody added. **(C)** CD4+ cells were added to HCMV-infected autologous fibroblasts at 1 dpi either in isolation, with anti-MHC Class II antibody, anti-IFN-γ antibody, or isotype to anti-MHC Class II antibody and incubated for a further 7 days before harvest and analysis by flow cytometry. Error bars represent SD of triplicates. One experimental replicate out of 3 is shown. *p < 0.05, **p < 0.01, ***p < 0.001.

IFN-γ exerts its antiviral effects via a number of mechanisms, which includes induction and upregulation of MHC Class II expression.^39^ We hypothesised that IFN-γ induction of MHC Class II was essential to allow CD4+ cells to mediate antiviral activity and therefore neutralization of IFN-γ or blocking MHC Class II antigen presentation would inhibit the antiviral activity. CD4+ cells were isolated by MACS using positive selection beads. They were then added to the HCMV-infected fibroblasts either alone, with anti-MHC class II antibody or anti-IFN-γ antibody, or isotype control antibody at E:T ratios of 2.5:1, 1.25:1, 0.6:1 and 0.3:1 and co-cultured for a further 7 days. The results showed that antibody neutralisation of IFN-γ reduced anti-viral activity of CD4+ cells (seen as increased IE and late CMV gene expression) particularly at the lower E:T ratios of 1.25, 0.6 and 0.3:1. (Fig.4C, teal bars). Also, the addition of blocking anti-MHC class II antibodies (Fig.4C, pink bars) was able to reduced anti-viral activity of the CD4+ cells. It was noted that at higher E:T ratios, the percentage of inhibition by the blocking antibodies decreased.

In summary, these experiments showed that addition of CD4+ cells induced the expression of MHC Class II on the cell surface of uninfected fibroblasts, which allowed antigen presentation to CD4+ T cells via the MHC Class II pathway when these cells became infected, leading to the antiviral effect of CD4+ cells. In addition, these results also showed that induction of these MHC Class II complex was mediated by IFN-γ.

### CD4+ T cells require a population of monocytes in order to limit HCMV gene expression

Our experiments so far showed that CD4+ cells from HCMV-seropositive donors co-cultured with HCMV-infected autologous fibroblasts were able to control viral gene expression and spread. They also showed that this antiviral effect was mediated by IFN-γ and by induction of MHC Class II on the cell surface. However, this still did not explain how CD4+ cells added to the viral dissemination assay were able to initially recognise HCMV-infected fibroblasts, in the absence of constitutive fibroblast expression of MHC Class II.

CD4 is predominantly expressed by CD4+ T cells, but also by a minor population of CD4+CD8+ T cells, monocytes subsets,^40^ dendritic cell subsets (both plasmacytoid DCs and conventional/myeloid DCs),^41,42^ and a small number of neutrophils,^43^ CD34+ progenitor cells,^44,45^ and NK cells.^46^ All previous experiments were performed using CD4 cells isolated by positive selection, however it is likely that this approach could co-isolate other cells that might express MHC Class II and have antigen presentation capacity, such as myeloid and dendritic cells. The presence of MHC Class II expressing antigen presenting cells might enable the initial HCMV antigen presentation to CD4+ T cells, which could trigger IFN-γ production and the subsequent MHC Class II induction on fibroblast which are then able to themselves able present antigen when they become HCMV infected.

Consequently, we tested this hypothesis by isolating CD4+ cells using two different methods: a direct CD4 Microbead kit (Miltenyi), which would select all cells that express CD4, and a CD4+ T cell Isolation kit (Miltenyi) which isolated “pure” CD4+ T cells by removing cells which expressed CD8, CD14, CD15, CD16, CD19, CD36, CD56, CD123, TCR γ/δ, and CD235a. These cell preparations were stained with anti-CD45, CD3, CD4 (to identify CD4+ T cells), CD14, CD16, (to identify monocyte/myeloid sub-populations) and CD11c, CD123, CD303 and HLA-DR (to identify DC sub-populations). The results clearly show that a population of CD3-CD4+ cells were present in the cells isolated using direct anti-CD4 Microbeads (Fig.5A, red outline). Gating on this population of cells using the other phenotyping markers demonstrated that these CD3-CD4+ cells contained populations of classical, intermediate and non-classical monocytes. In addition, among the CD14-CD16-population of cells, there was a population of cells which were CD11c-CD303+ and intermediate for HLA-DR and CD123, and another population which was CD11c+CD303-with higher levels of HLA-DR and CD123 (Fig.6A). Therefore, these cells were likely to be plasmacytoid dendritic cells (pDCs) and conventional dendritic cells (cDCs), respectively. In contrast, this population of CD3-CD4+ cells were absent in the cells selected using the CD4+ T cell Isolation Kit (Fig.6B, green outline).

**Figure 5.**
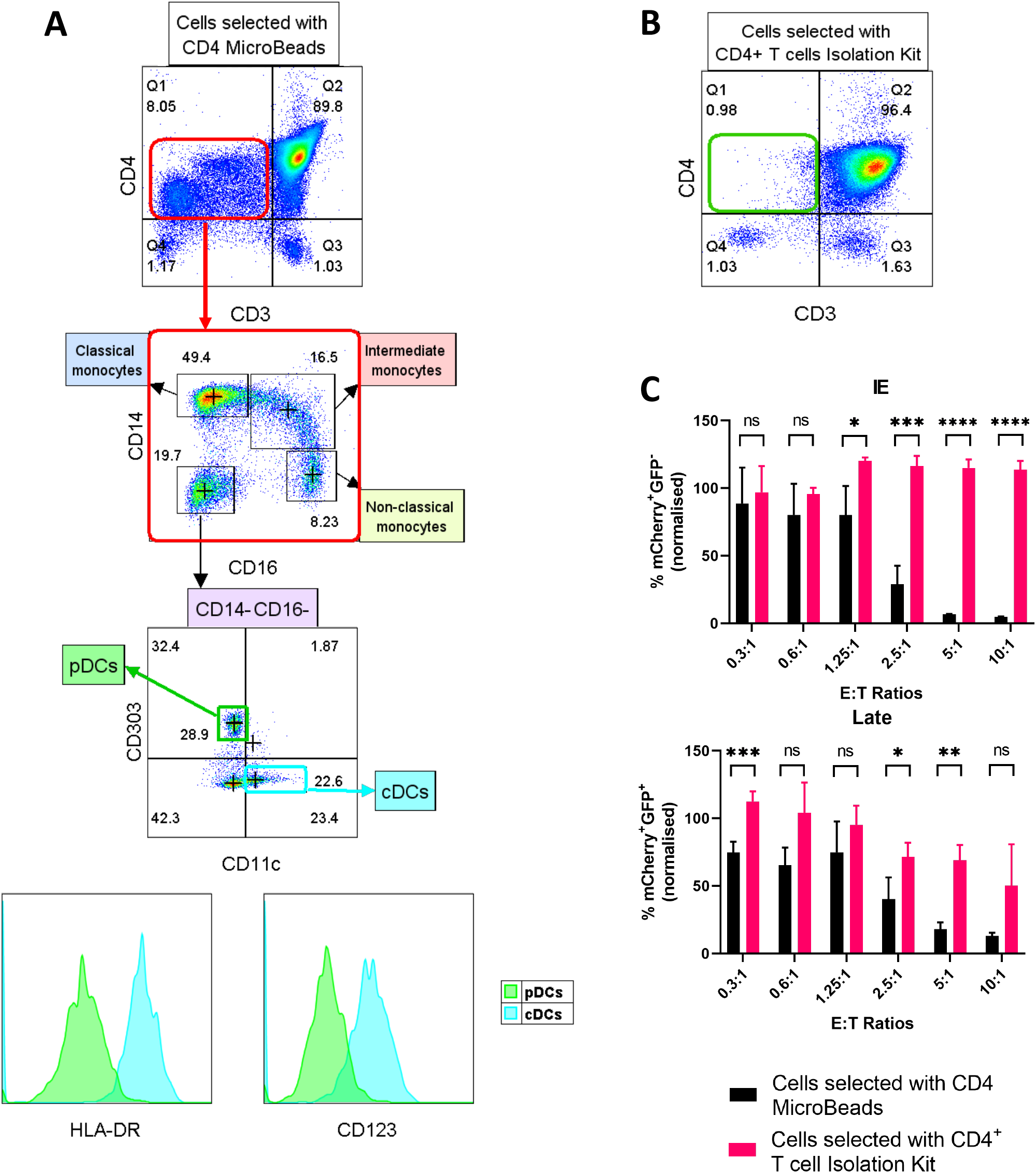
CD4+ T cells in isolation do not control viral dissemination, but cells selected with anti-CD4 Microbeads do. PBMCs from an HCMV-seropositive donor underwent positive selection with anti-CD4 Microbeads **(A)** or negative selection with the CD4 T cell Isolation Kit **(B**) and stained with antibodies to CD45, CD3, CD4, CD14, CD16, CD11c and CD303. There was a population of CD3-CD4+ cells in the cells which underwent positive selection (red outline), which was absent in the cells which underwent negative selection (green outline). These cells were found to consist of populations of classical, intermediate and non-classical monocytes, and CD14-CD16-cells. The CD14-CD16-population was found to contain populations of plasmacytoid dendritic cells (pDCs) and conventional dendritic cells (cDCs). **(C)** Primary dermal fibroblasts from an HCMV-seropositive donor were infected with HCMV (MOI = 0.27). At 1 dpi, autologous PBMCs underwent magnetic column sorting with either anti-CD4 Microbeads or CD4+ T Cell Isolation Kit and the resultant immune cell populations were added to the HCMV-infected primary dermal fibroblasts. Plates were incubated for a further 7 days before harvest and analysis by flow cytometry. Error bars represent SD of triplicate wells, statistics performed using Student’s *t*-test. *p < 0.05, **p < 0.01, ***p < 0.001, ****p < 0.0001.

**Figure 6.**
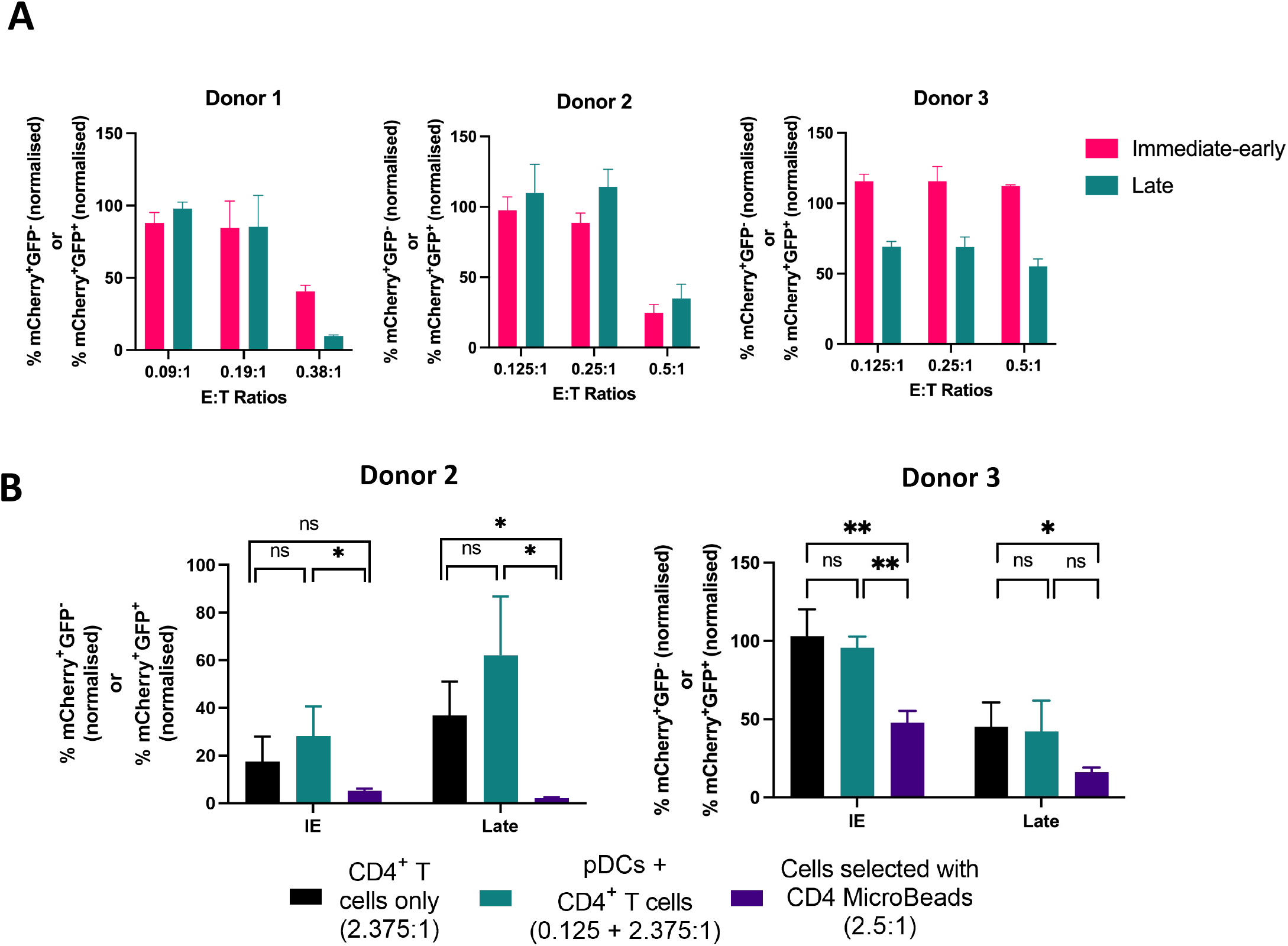
Co-culture of CD4+ T cells with plasmacytoid DCs does not contribute to control of IE and late CMV gene expression. **(A)** Plasmacytoid dendritic cells from 3 donors were added to HCMV-infected autologous fibroblasts at 1 dpi followed by incubation for 7 days before harvest and analysis by flow cytometry. **(B)** CD4+ T cells obtained by depletion from 2 HCMV-seropositive donors were added either in isolation or with plasmacytoid dendritic cells from the same donor to autologous HCMV-infected fibroblasts at 1 dpi. CD4+ cells obtained by positive selection with anti-CD4 MicroBeads were also added to HCMV-infected autologous fibroblasts in separate wells. The E:T ratios at which these cells were added is shown in the legend. These wells underwent a further 7 days’ incubation before harvest and analysis by flow cytometry. Error bars represent SD of triplicate wells, statistics performed using Student’s *t*-test. *p < 0.05, **p < 0.01.

We then questioned if the “pure” CD4+ T cells isolated using the CD4+ T cell Isolation Kit would show the same antiviral effect as CD4+ cells isolated using the CD4 Microbeads kit. To investigate this, these two cell preparations were used separately in the VDA over a range of E:T ratios. As expected, directly isolated CD4+ cells were antiviral, inhibiting both IE and late gene expression. However, in contrast, the “pure” CD4+ T cells could not suppress IE expression and only demonstrated inhibition of late gene expression at high E:T ratios. (Fig.6B) Taken together, these results suggest that CD4+ T cells require the presence of either the monocyte and/or the DC populations in order to drive CD4+ T cell-mediated virus control. Therefore, in order to determine the contribution of these populations, we isolated purified CD4+ T cells as well as the CD14+ myeloid fraction and pDC sub-populations to perform co-culture experiments.

### Co-culture of CD4+ T cells with plasmacytoid dendritic cells does not contribute to control of IE and late HCMV gene expression

Monocytes and some dendritic cells constitutively express MHC Class II on their cell surface and thus do not require IFN-γ to induce expression of MHC Class II for antigen presentation to CD4+ T cells.^47^ The presence of these cell populations could allow antigen presentation via the MHC Class II pathway to occur, and provide a possible explanation for how CD4+ cells from seropositive donors were able to recognise HCMV-infected fibroblasts in the assay.

We first examined if pDCs added to HCMV-infected autologous fibroblasts were antiviral. pDCs from 3 donors were added to infected autologous fibroblasts (MOI = 0.03) at 1 dpi at E:T ratios of 0.38 to 0.5:1 and incubated for a further 7 days before harvest and analysis by flow cytometry. The results from donor 1 and 2 showed that there was an inhibitory effect of plasmacytoid dendritic cells on immediate-early and late HCMV gene expression at the highest E:T ratio tested, but that this effect rapidly diluted out. The results from donor 3 showed that there was an inhibitory effect on HCMV late gene expression at the E:T ratio of 0.5:1, but that there was no inhibitory effect on HCMV immediate-early gene expression (Fig.6A).

From our phenotyping experiments, we expected that pDCs made up approximately 1.5-2% of the non-granulocyte population of cells obtained by anti-CD4 Microbeads. In an experiment where cells selected with anti-CD4 Microbeads were added to infected fibroblasts at an E:T ratio of 10:1, this would be equivalent to an E:T ratio of 0.15-0.2:1 of pDCs. At a starting pDC E:T ratios of 0.38 to 0.5:1, which was approximately 2.5 fold higher than present in the pDC fraction present in the direct CD4 isolated cells, an antiviral effect of pDCs was observed. However, this antiviral effect diluted out rapidly as the E:T ratios decreased. We thus concluded that It was therefore unlikely that pDCs contributed to a direct antiviral effect in the viral dissemination assay when used at the E:T ratios in our earlier experiments.

We then proceeded to examine if pDCs participated in antigen presentation to CD4+ T cells in the viral dissemination assay and contributed to the antiviral effect of CD4+ T cells on HCMV-infected fibroblasts. Using directly isolated CD4 cells as a positive control, “pure” CD4+ T cells from a seropositive donor were added to HCMV-infected autologous fibroblasts at 1 dpi either in the presence or absence of autologous pDCs. Where both “pure” CD4+ T cells and pDCs were added together, the ratio of CD4+ T cells to pDCs was 2.375:0.125 (i.e. 95% of immune cells added were CD4+ T cells, 5% were pDCs). The results showed that, while CD4+ cells obtained using anti-CD4 Microbeads had an antiviral effect as we have previously shown, “pure” CD4+ T cells had minimal or significantly reduced antiviral activity, also as expected. However, addition of pDCs at a ratio slightly higher than would be present in the total CD4+ cell population did not restore the antiviral activity (Fig. 6B). Taken together, the results suggest that it was unlikely that the pDCs contributed to driving the antiviral function of the CD4+ T cells.

### CD14+ monocytes contributes to control of IE and late CMV gene expression by CD4+ T cells, which is mediated by IFN-γ and MHC Class II

The other major population of cells which was present in preparations obtained by anti-CD4 Microbeads were monocytes. We therefore used the same approach as for pDCs to determine if co-culture of monocytes in the viral dissemination assay restored the antiviral function of CD4+ T cells. “Pure” CD4+ T cells obtained by depletion and CD14+ monocytes obtained by positive selection using CD14 Microbeads were added to HCMV-infected autologous fibroblasts (MOI = 0.03) at 1 dpi, either in isolation or together. The CD4+ T cells were added at an E:T ratio of 9:1 and the CD14+ monocytes were added at an E:T ratio of 1:1. As a positive control, CD4+ cells obtained by positive selection using anti-CD4 beads were used, added at an E:T ratio of 10:1.

The results show that “pure” CD4+ T cells and CD14+ monocytes had little antiviral effect when added in isolation. However, when co-cultured together, they restored the antiviral activity of the CD4+ T cells to a level similar to that seen when CD4+ cells obtained using anti-CD4 Microbeads were added to the viral dissemination assay (Fig.7A). Fluorescence microscopy images of the cells also showed the presence of mCherry fluorescence in the immune cells of the wells which had monocytes added, but not in the wells which had “pure” CD4+ T cells added, suggesting that that the CD14+ monocytes had taken up viral antigen (Figs.7B—D).

**Figure 7.**
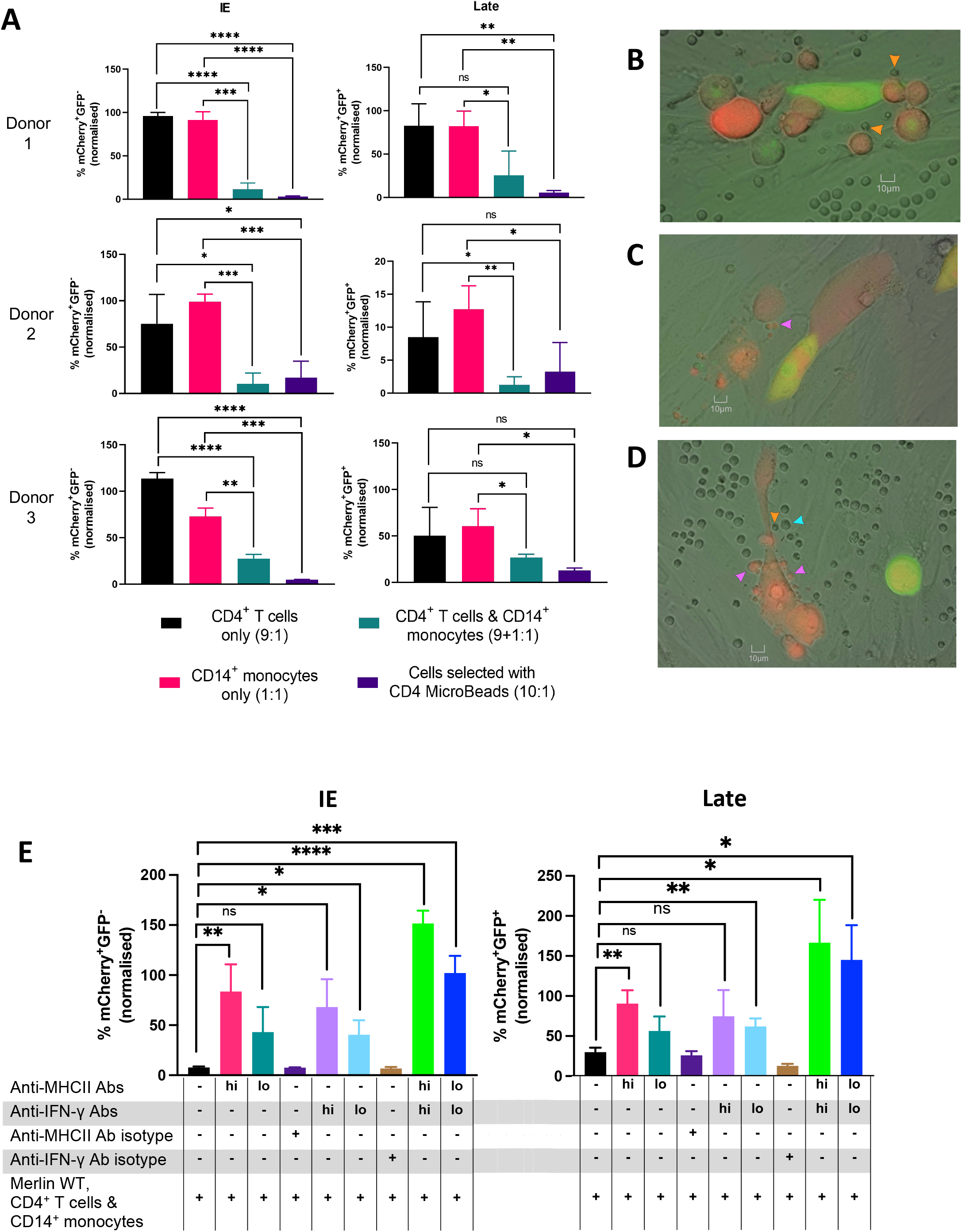
Co-culture of CD4+ T cells with CD14+ monocytes potentiates control of IE and late CMV gene expression, which is mediated by IFN-γ and MHC Class II. **(A)** CD14+ monocytes obtained by positive selection and CD4+ T cells obtained by depletion were obtained from PBMCs of 3 HCMV-seropositive donor by magnetic column sorting and added to HCMV-infected autologous fibroblasts at 1 dpi either in isolation, or together, followed by incubation for a further 7 days before harvest and analysis by flow cytometry. The E:T ratios at which these cells were added is shown in the legend. Microscopy images of **(B)** CD4+ T cells only, **(C)** CD14+ monocytes only, **(D)** CD4+ T cells and CD14+ monocytes together. (Orange arrowheads = monocytes that have taken up fluorescence, purple arrowheads = CD4+ T cells, cyan arrowhead = monocytes that have not taken up fluorescence.) **(E)** CD4+ T cells and CD14+ monocytes were added (in a proportion of 9:1) to HCMV-infected autologous fibroblasts at 1 dpi with anti-MHC Class II antibody or anti-IFN-γ antibody at either high doses (50*μ*g/ml for anti-MHCII; 10*μ*g/ml for anti-IFN-γ; indicated by “hi”) or low doses (10*μ*g/ml for anti-MHCII; 3*μ*g/ml for anti-IFN-γ; indicated by “lo”) and incubated for a further 7 days before harvest and analysis by flow cytometry. Error bars represent SD of triplicate wells, statistical analysis performed with Student’s *t*-test. *p < 0.05, **p < 0.01, ***p < 0.001, ****p < 0.0001.

Finally, we then sought to determine if these monocytes restored the CD4+ T cell control of HCMV-infected fibroblasts by antigen presentation via the MHC Class II antigen presentation pathway. “Pure” CD4+ T cells and CD14+ monocytes were added to HCMV-infected autologous fibroblasts (in a proportion of 9:1), at an E:T ratio of 1.25:1, at 1 dpi. At the same time that the immune cells were added, anti-MHC Class II antibody or anti-IFN-γ antibody was also added, either in isolation or together, and at either high doses (50*μ*g/ml for anti-MHCII; 10*μ*g/ml for anti-IFN-γ) or low doses (10*μ*g/ml for anti-MHCII; 3*μ*g/ml for anti-IFN-γ). The cells were incubated for a further 7 days before harvest and analysis by flow cytometry. The results demonstrated that blocking MHC Class II or neutralizing IFN-γ antibodies at the higher dose separately showed a partial abrogation of the antiviral effect (Fig.7E, pink and lilac bars). However, in combination, this blocked antiviral activity completely (Fig.7E, green bars). In addition, this effect was dose dependent (Fig.7E, teal, light blue and dark blue bars).

Taken together, these results suggest that these CD14+ monocytes are able to take up HCMV antigens and present them to HCMV-specific CD4+ T cells via the MHC Class II antigen presentation pathway. This then results in the production of IFN-γ, which induces expression of MHC Class II on fibroblasts that had not yet progressed to the immediate-early phase of infection. Subsequently, when these cells are in the immediate-early phase of infection, this then allows continued antigen presentation to HCMV-specific CD4+ T cells via MHCII, leading to persisting antiviral cytokine production and control of viral spread in the fibroblasts by these HCMV-specific CD4+ T cells.

## DISCUSSION

While the role and mechanisms of HCMV specific CD8+ T cell control of HCMV has been extensively investigated, the function of HCMV specific CD4+ T cells is less well characterized. Yet there are multiple clinical observation of transplant patients with poorer CD4+ T cell recovery being at higher risk of CMV viraemia and disease.^16–19^ While HCMV-specific CD4+ T cells could provide help to HCMV specific CD8+ T cells in order to exert antiviral activity they also have a direct antiviral role themselves through production of antiviral cytokines and direct cytolytic activity.^48^ The results presented in this paper investigate a mechanism by which HCMV-specific CD4+ T cells could recognise HCMV-infected cells that do not constitutively express MHC Class II, which would allow these CD4 T cells to directly exert their antiviral effect. The results also highlight the essential contribution of monocytes in enabling induction of a CD4+ T cell IFN-γ response, which would stimulate MHC Class II expression on cells that do not normally express these molecules and facilitate an amplification of HCMV-specific CD4+ T cell activation and direct anti-viral CD4+ T cell effector functions, such as cytotoxicity.

Many of the cells infected by HCMV during end-organ disease do not constitutively express MHC Class II. Recognition of these infected cells by HCMV-specific CD4+ T cells would require induction of MHC Class II complexes on their cell surface, which can be induced by IFN-γ via the Class II transcriptional activator (CIITA) gene. This paper has made extensive use of autologous primary dermal fibroblasts as these cells will support full lytic and spreading infection of HCMV. These cells are readily derived from a relatively simple small punch biopsy, and critically they express the correct MHC Class I and II alleles essential for T cell recognition of antigen presentation. This gives a tractable *in vitro* model to investigate the ability of lymphocyte subsets to inhibit viral spread and replication.

Fibroblasts are an important target cell for the replication of virus *in vivo*. However, other cell types such as epithelial cells, endothelial cells, smooth muscle cells as well as DCs and macrophages are also infected by HCMV *in vivo*. Infection of these cells contributes to *in vivo* dissemination and cell damage leading to end organ disease,^49^ but deriving autologous epithelial and endothelial cells for these studies is not feasible. It is possible that the immune recognition of these other cell types when infected with HCMV is different from the results obtained studying dermal fibroblasts.

However, we have demonstrated that epithelial and endothelial cells that also do not constitutively express MHC Class II can be induced to express it when exposed to secretomes from CD4 cells co-cultured with HCMV-infected fibroblasts (Supp Fig.5). As such, it is likely that these cells could also be recognized by HCMV-specific CD4 T cells.

We have demonstrated that the supernatants of CD4+ cells co-cultured with HCMV-infected autologous fibroblasts were antiviral, and IFN-γ was found to be the highest-upregulated cytokine in these secretomes. IFN-γ is an antiviral cytokine produced by activated CD4+ T cells. Upon binding to its receptor, there is activation of IFN-γ-stimulated gene transcription, which leads to induction of an antiviral state via transcription of antiviral genes.^37^ Binding of IFN-γ to its receptor also triggers induction of MHC Class II on the cell surface via Class II transactivator gene transcription.^50^ Antibody-mediated neutralization of IFN-γ completely abrogated MHC Class II induction and as such is both necessary and sufficient. These effects of IFN-γ are also seen on uninfected and HCMV-infected fibroblasts (Supp Fig.4). The secretome also contains a number of other classic antiviral factors at lower levels than IFN-γ—namely, type I and III interferons as well as TNF-α. It was however notable that these secretomes could inhibit both viral spread and late gene expression, but when washed off late virus gene expression and subsequent virus was observed—suggesting that this was a virustatic and not a virucidal effect.

The presence of two other cytokines in the CD4 secretome were also notable—GM-CSF and IL-10. GM-CSF is secreted by activated macrophages, lymphocytes (NK and T cells), as well as fibroblasts and endothelial cells.^51^ It is a polyfunctional cytokine which drives tissue inflammation and is a growth factor, and signalling through its receptor activates an number of key signalling pathways including MAPK, JAK/STAT, NFkB and PI3K.^52^ It drives the activation of myeloid cells and the secretion of antiviral cytokines such as TNFa and IL-1b both present in the CD4 secretomes.^53^ Taken together, this can help drive an antiviral state in surrounding cells prior to HCMV infection.^54^

IL-10 on the other hand is a an anti-inflammatory immunosuppressive cytokine which acts to inhibit typical Th1 cytokines such as IFN-γ as well as pro-inflammatory cytokines TNF-α and GM-CSF.^55^ It has been shown that a number of HCMV-specific CD4+ T cells, especially those specific for viral proteins that are expressed during both lytic and latent infection, secrete IL-10.^56^ This effect has been shown to produce a suppressive environment that might protect HCMV latently infected cells from being recognised and eliminated by T cells but might also be modulating T cells responses in lytic infection. It is clear in this assay system where virus is spreading in the presence of HCMV specific memory T cells that the overall effect is overall control of virus infection but not its elimination.

We have also demonstrated that co-culture of CD4+ cells with virus-infected cells controls virus dissemination and late gene expression, and induces MHC II expression on autologous fibroblasts. We have also shown that this is blocked by either anti-IFN-γ or anti-MHC II antibodies within the context of the VDA. The results suggest that MHC Class II presentation is required to drive the antiviral effect and that if IFN-γ is removed the antiviral effect is also blocked (Fig 4). It was however clear that CD4+ T cells prepared using an isolation technique based on negative selection were unable to mediate anti-viral activity. Given that fibroblasts do not constitutively express MHC Class II we hypothesised that there must be an initial presentation of HCMV antigens via MHC Class II-expressing cells to HCMV-specific T cells. This would allow the production of IFN-γ from these T cells and subsequent induction of MHC Class II on the fibroblasts that do not constitutively express it. We subsequently demonstrated that the presence of myeloid cells, and in particular CD14+ monocytes in the total CD4+ cell co-cultures, were essential in order to drive control of HCMV in these assays (Figs 6 and 7).

We did not assess the contribution of conventional dendritic cells to CD4+ T cell control, or overall control, of HCMV-infected cells. As the majority of the CD3-CD4+ population in the cells obtained by anti-CD4 Microbeads were monocytes, it would appear that the percentage of conventional dendritic cells in this population was likely to be less than 10% of these cells, therefore the overall contribution of this cell type was likely to be low. Prior work by our group has shown that conventional dendritic cells can be lytically infected with HCMV, and that co-culture of these cells with CD4+ cells leads to control of viral gene expression.^27^ Thus, we have not repeated those experiments here and have focused on CD14+ monocytes and pDCs instead.

It is also interesting to note that adaptive NK cells (expressing NKG2C) also constitutively express HLA-DR, and that these NK cells can take up HCMV antigens which they are then able to present antigen to HCMV specific CD4+ T cells.^57^ This NK cell-mediated antigen presentation resulted in IFN-γ and TNF-α secretion, as well as the expression of CD107a, an indicator of cytotoxic degranulation. As such, we would speculate that HCMV replication and inflammation that recruited NK cells could further enhance activation of CD4+ T cells by production of IFN-γ, inducing MHC Class II on multiple cells that do not constitutively express MHC Class II.

It has been recognised for some time that HCMV generates Th1 CD4+ T cells that secrete a number of antiviral cytokines (IFN-γ and TNF-α) which are also directly cytotoxic in an MHC Class II-dependent fashion. Direct cytolytic activity via both perforin- and Fas-dependent killing mechanisms has been described.^25,27,48,56,58–61^ HCMV-specific cytotoxic T cells have also been described to emerge following CMV infection^26,62^ and have demonstrated a capability to lyse CMV antigen-expressing target cells *in vitro*.^25^ This suggests that CMV lytically-infected, MHC Class II-expressing cells could be targeted by these T cells in vivo. It is tempting to speculate if this is why a lack of CD4 cells has been associated with viraemia in a number of transplant scenarios.^16,63,64^

It is well recognised that HCMV encodes multiple immune evasion proteins which downregulate MHC Class II complexes on the cell surface. US2 binds to MHC Class II-α chains and assembled MHC Class II-α/β/I_i_ complexes, leading to their degradation,^65^ while US3 alters assembly of MHC class II complexes by binding HLA-DR proteins before or during assembly of α/β complexes in the endoplasmic reticulum, preventing the binding of the invariant chain. This leads to mislocalisation of these complexes to other post-Golgi compartments and results in the reduction of antigen presentation in US3-expressing cells.^66^ UL23 interferes with IFN-γ-mediated MHC Class II gene transcription, and UL31 and pp71 inhibit IFN-associated gene transcription.^65–71^ However, as the CD4+ T cells in our *in vitro* assay were able to exhibit an antiviral effect, we believe that IFN-γ establishes an environment whereby subsequent infection of MHC Class II-expressing fibroblasts (viral dissemination) leads to presentation of viral antigen to HCMV-specific CD4+ T cells before downregulation of MHC Class II is mediated by HCMV immune evasion genes, thus creating a window of opportunity to recognise infected cells early post infection.

In conclusion, we have shown in this paper that the antiviral activity of CD4+ T cells against HCMV-infected autologous fibroblasts is mediated by antigen presentation from CD14+ monocytes via the MHC Class II pathway. However, HCMV is able to lytically infect a wide variety of human cells including fibroblasts, epithelial cells and endothelial cells, smooth muscle cells as well as DCs and macrophages both *in vivo* and *in vitro*.^49^ *In vivo*, infection of these cells contributes to viral dissemination and cell damage, ultimately leading to end organ disease. It is essential to be able to derive autologous HLA Class I and II-mapped cells in order to be able to study T cell immune recognition and control of HCMV infected cells. To date, we have been limited to using autologous fibroblasts as these can be readily derived from minimally invasive dermal punch biopsies and grown out in primary cell culture. While fibroblasts are an important target cell for the replication of virus *in vivo*, it is possible that the immune recognition of these other cell types when infected with HCMV is different from the results obtained studying dermal fibroblasts. With the advent of de-differentiation technologies, it is now feasible to use established primary fibroblast cell cultures as the starting point to derive human induced Pluripotent Stem Cells (hiPSC) from them. These hiPSC cells can then be re-differentiated into other cell types—in particular, endothelial cells and myeloid cells or DCs. Importantly, these cells will be autologous to the original donor and can then be used to determine cell-mediated control of HCMV in these clinically relevant targets for *in vivo* HCMV replication.

## MATERIALS AND METHODS

### Recruitment of healthy donors and fibroblasts

HCMV seronegative and seropositive donors were recruited locally with ethical approval from the Cambridge Central Research Ethics Committee (97/092) or via the NIHR BioResource Centre Cambridge through the AQUARIA study, with ethical approval from the North of Scotland Research Ethic Committee (NS/17/0110). Donors were excluded if they were receiving immunosuppressive therapy, e.g., cyclosporins or methotrexate. In each case, informed written consent was obtained from all donors in accordance with the Declaration of Helskinki.

Primary dermal fibroblasts and PBMCs were obtained from these donors. HCMV serostatus was assess using an IgG enzyme-linked immunosorbent assay (Trinity Biotech Plc, Co., Ireland). Fibroblasts were obtained by a 2-mm punch biopsy followed by growing out into a fibroblast cell line using a previously published protocol^72^ modified to use DMEM^73^. PBMCs were isolated from heparinised blood samples using Histopaque®-1077 (Sigma-Aldrich, UK) or Lymphoprep (Axis-shield, Norway) density gradient centrifugation. PBMCs were frozen in liquid nitrogen in a 10% dimethyl sulfoxide (DMSO) (Sigma-Aldrich, UK) and 90% SeraPlus foetal bovine serum (FBS) (PAN Biotech, UK) solution or serum-free freezing media composed of 60% Isocove’s modified Dulbecco’s Medium, (Sigma-Aldrich, UK), 10% DMSO and 30% Panexin serum replacement (PAN Biotech). When required for use, frozen PBMCs were rapidly thawed by hand warming and diluted into 10mls of X-Vivo 15 (Lonza, UK) followed by centrifugation and resuspension in fresh X-Vivo 15 to remove DMSO. PBMCs were incubated at 37°C for 1 hour with 10U/ml DNase (Benzonase, Merck-Millipore via Sigma-Aldrich) followed by centrifugation, resuspension in fresh media and overnight rest at 37°C before use.

### Isolation of cells using magnetic columns

Immune cell populations were obtained from PBMCs using magnetic-activated cell sorting by autoMACS Pro Separator® (Miltenyi, U.K.) in accordance with manufacturer’s instructions. CD4+ and CD14+ cells were selected by positive selection using CD4 and CD14 Microbeads respectively, and CD4+ T cells were selected by depletion of unwanted cell populations using the CD4+ T Cell Isolation Kit (all reagents from Miltenyi, UK).

### Fluorescently labelled Merlin HCMV (mCherry-P2A-UL36 [vICA], GFP-UL32 [pp150])

The virus used in this study was based on a BAC cloned version of HCMV strain Merlin^74^. This contains the complete wildtype HCMV genome, with the exception of point mutations in RL13 and UL128, which enhance growth in fibroblasts^75^. Two genes were tagged with fluorescent markers. UL36 was linked to mCherry via a peptide-2A linker, which results in expression of mCherry at immediate-early times post-infection^76^, and UL32 was linked directly to GFP via a six amino-acid linker, which results in GFP expression at late times^77^. Both constructs were generated by recombineering as previously described^78^.

### Secretome generation

To generate the PBMC or CD4+ cell secretomes, CD4+ cells were isolated from PBMCs by automated magnetic activated cells sorting (autoMACS, Miltenyi, U.K.) using CD4 MicroBeads (Miltenyi, U.K.) as described above. Following purity analysis, the PBMC and CD4+ cell fractions were then added to either HCMV-infected or uninfected autologous primary dermal fibroblasts at E:T ratios of 10:1 in 24-well plates in Roswell Park Memorial Institute (RPMI)-1640 media (Sigma-Aldrich, UK) supplemented with 10% FBS and 100 units/ml of penicillin/streptomycin. After one week, media was harvested from the wells and supernatant from the wells obtained by centrifugation at 935G for 10 minutes and stored at -80°C until needed.

### Cell culture and viral dissemination assay

Human foetal foreskin fibroblasts, primary dermal fibroblasts and ARPE-19 cells were grown and maintained in 175cm^2^ tissue culture flasks (Corning, U.K.) and maintained in Dulbecco’s Modified Eagle Medium (DMEM) (high glucose) (Sigma-Aldrich, U.K.) supplemented with 10% foetal bovine serum (FBS) and 100 units/ml of penicillin/streptomycin. Human umbilical vein epithelial cells (HUVECs) were a gift from Jing Garland and grown in 75cm^2^ tissue culture flasks (Corning, U.K.) and maintained with Endothelial Cell Growth Medium 2 supplemented with Endothelial Cell Growth Medium 2 Supplement Pack (PromoCell, Germany).

For the viral growth and dissemination assays, fibroblasts were seeded in 96-well full-area plates (Corning, U.K.) at 20,000 cells per well, or 96-well half-area plates (Greiner Bio-One, USA) at a density of 10,000 cells per well and grown to confluency overnight in DMEM at 37°C, 5% CO_2_. When cells were confluent, infections were performed by adding virus diluted in same medium as the cells to achieve the required multiplicity of infection. At the end of the incubation periods, cells were harvested with trypsin and fixed in a 2% paraformaldehyde solution for flow cytometry analysis.

The data was then plotted as graphs showing amount of infection on the *y*-axis, which was either expressed as a percentage of total cells (“% mCherry+GFP-”) or as a percentage of the amount of cells in the same phase of CMV gene expression in the infected controls of the experiment, i.e. “normalised to infected controls” (“% mCherry+GFP- (normalised)” or “% mCherry+GFP+ (normalised)”). The equation below gives an example of the calculation for % mCherry+GFP- (normalised).

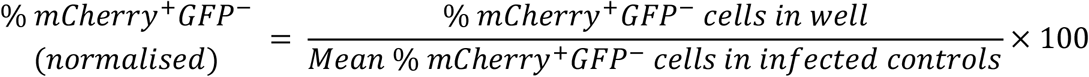

### Cellular staining (IE72) for fluorescence microscopy

Cells were fixed with 1% paraformaldehyde for 15 minutes and permeabilised with 70% ethanol at −20°C for 20 minutes before incubation with 5% milk for 1 hour at 22–26°C, before milk was removed and E13 antibody (bioM’erieux, U.S.A.) was added at 1:1000 dilution for 1 hour. This was followed by 3 washings with TBS Tween, before incubation for 1 hour with secondary antibody (goat anti-mouse IgG1 conjugated to AlexaFluor 788) at 1:500 dilution with 5% milk. This was followed by two washings with TBS Tween. PBS was then added to cells and fluorescence microscopy carried out with a widefield Nikon TE200 microscope with CoolLED pE-4000 as UV light source, and digital images taken with Image-Pro Premier 9.3 software. Images were processed using ImageJ (available at https://imagej.nih.gov/ij/).

### Real-time quantitative Polymerase Chain Reaction

RNA extraction was performed with RNEasy Kit using RNEasy spin columns (both from QIAgen, U.K.) according to manufacturer’s instructions. Removal of genomic DNA and subsequent complementary DNA was made using QuantiTect Reverse Transcription Kit (QIAgen, U.K.) as per manufacturer’s instructions. RT-qPCR analysis was performed using New England Biotech LUNA SYBR Green qPCR reagents (New England Biolabs, U.K.). Glyceraldehyde phosphate dehydrogenase (GAPDH) was used as reference genes and relative gene expression was analysed using 2∧ΔCt values (cycle threshold values). The ΔCt value was calculated using the equation below: ΔCt = Ct value of gene of interest – GAPDH Ct value. Primer sequences were: UL44 (Forward) TACAACAGCGTGTCGTGCTCCG, (Reverse) GGCGTAAAAAACATGCGTATCAAC; pp28/UL99 (Forward) TTCACAACGTCCACCCACC, (Reverse) GTGTCCCATTCCCGACTCG.

### Cellular surface staining, phenotyping and post-separation purity analysis

A minimum of 10^5^ of the cells of interest were stained using the following antibody-fluorochrome combinations: CD3-VioGreen, clone BW264/56 (Miltenyi Biotec); CD4-Brilliant Violet 605, clone OKT4 (BioLegend); CD11c-APC, clone 3.9 (BioLegend); CD14-BV570, clone M5E2 (BioLegend); CD16-BV650, clone 3G8 (BioLegend); CD45-VioBlue, clone 5B1 (Miltenyi Biotec); CD123-PerCP/Cy5.5, clone 6H6 (BioLegend); CD303-FITC, clone 201A (BioLegend); HLA-DR-PE-Cy5 or HLA-DR-BV421, clone L243 (BioLegend).

Following magnetic column sorting, the purities of the cell populations obtained were assessed by cell surface staining of markers of interest using the following antibody-fluorochrome combinations: CD3-AF647, clone HIT3a (BioLegend) or CD3-FITC, clone UCHT1 (BioLegend); CD4-PE, clone RPA-T4 (BioLegend); CD14-FITC, clone 61D3 (eBioScience); CD8a-PerCP/Cy5.5, clone HIT8a (BioLegend); Live-Dead Far Red (ThermoFisher Scientific).

### Neutralisation assays

Blocking antibodies and their isotypes used were as follows: anti-IFN-γ, polyclonal, Goat IgG (R&D Systems, USA), isotype: normal goat IgG control, polyclonal (R&D Systems, USA); anti-MHC Class II, mouse IgG2a, κ, clone Tu39 (BD BioSciences, UK), isotype: purified NA/LE mouse IgG2a, κ, clone G155-178 (BD Biosciences, UK).

### Quantification of amounts of cytokines in secretomes

Quantification of amounts of cytokines present in the secretomes was performed using the LegendPlex™ (BioLegend, UK) assay in accordance with manufacturer’s instructions.

### Cytokine array

For detection of relative amounts of cytokines in various supernatants, a Proteome Profiler™ Human XL Cytokine Array Kit (R&D Systems) was used in accordance with the manufacturer’s instructions. The chemiluminescent signal produced by each capture antibody was measured on x-ray film and average mean gray values from the duplicate of each analyte was obtained using using ImageJ. Then, the mean intensity of each analyte was calculated using the formula:

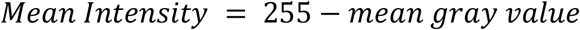

This was followed by subtraction of background mean intensity and then normalising the values by expression as a percentage of the mean intensity of standard reference spots provided on each membrane.

### Flow Cytometry

Flow cytometry analysis was performed on the BD Fortessa2 or Thermo Fisher Attune NxT flow cytometers, with assistance from the NIHR Phenotyping Hub at the University of Cambridge. Data was analysed with FlowJo v10 (Becton Dickinson, UK).

### FluoroSpot™ analysis

The FluoroSpot™ assay is an enzyme-linked immune absorbent spot (ELISpot)-based assay that allows quantitative measurement of frequencies of cytokine secretion of single cells. The assay was prepared as per manufacturer’s instructions. PBMCs of the donors to be analysed were defrosted, treated with DNase for 1 hour before washing off and rested overnight before undergoing magnetic-assisted cell separation to deplete CD8+ T cells. The immune cells were then resuspended in the appropriate volume of media to achieve a target of as close to 250,000 cells per 90μl of media as possible. 90μl of immune cells were then added to the wells, followed by the relevant peptides, mitogens (for positive control wells) or media (for negative control wells). 90μl of cells was also stained with an antibody mix containing antibodies to LiveDead FarRed, CD3, CD4 and CD8 before analysis with Accuri C6 flow cytometer to obtain anaccurate count of the number of CD3+ cells that had been added to the wells. The FluoroSpotTM plates were then incubated for 48 hours at 37°C, 5% CO_2_ before removal of media, washing and addition of tag-labelled secondary detection antibodies followed by fluorescently-labelled anti-tag antibodies, as per manufacturer’s instructions. The amount of IFN-γ secreted was then quantified by enumerating the number of fluorescence “spots” per well with a fluorescence reader, and given as the number of “spot-forming units” (sfu) per 10^6^ CD3+ cells added to the well.

### Gene Ontology analysis

Gene ontology analyses were performed using ClueGO version 2.5.9,^79^ a plug-in available on the Cytoscape platform (version 3.9.1).^80^

### Statistical and graphical analysis

Data presentation and statistical analyses were performed using GraphPad Prism v9.1.

## Acknowledgements

We would like to thank Dr Richard Stanton from Cardiff University who kindly provided the Merlin strain of HCMV virus used in these experiments. We also gratefully acknowledge the participation of all Cambridge NIHR BioResource volunteers, and thank the Cambridge BioResource staff for their help with volunteer recruitment. The Cambridge BioResource is funded by the National Institute for Health Research (NIHR) Cambridge Biomedical Research Centre (BRC) and the NHS Blood and Transplant (NHSBT). We also gratefully acknowledge the Cambridge NIHR BRC Cell Phenotyping Hub, in particular Veronika Romashova for her flow cytometry advice and support.

