## Supplemental Figures for "Antiviral activity of Cytomegalovirus-specific CD4+ T cells against lytically infected non-haemopoietic cells"

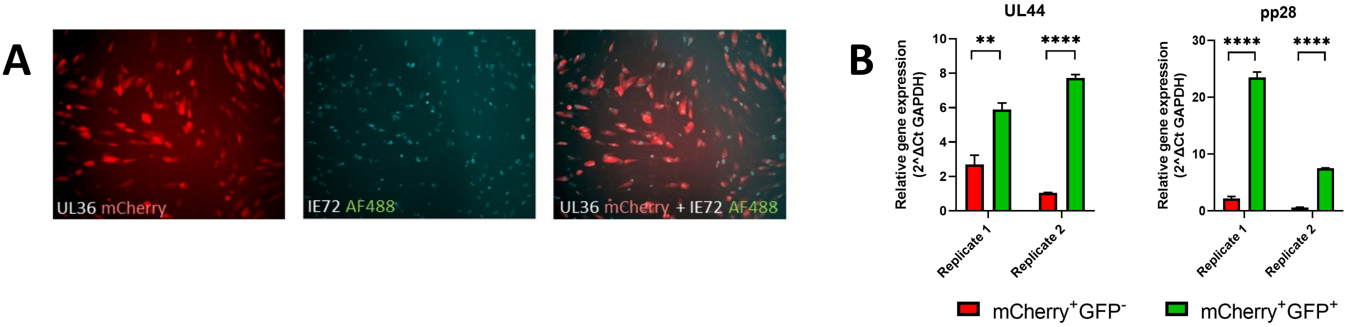


**Supp Fig 1. Cells expressing mCherry are in immediate-early phase of HCMV gene expression, and those expressing GFP are in late phase**

**(A)** Primary dermal fibroblasts were infected (MOI=0.03) with a dual-fluorescently tagged HCMV strain (Merlin mCherry-P2A-UL36 [vICA], GFP-UL32 [pp150]) which expresses mCherry when the virus enters the immediate-early life cycle, and GFP when the virus enters the late life cycle. Flow cytometry is used to quantify the percentage of fibroblasts which are in IE viral gene expression (mCherry+GFP-) and late gene expression (mCherry+GFP+). At 20 hours post-infection, cells were stained for IE72. Left image shows microscopy images of cells expressing mCherry, middle image shows same cells stained for IE72. Right image shows superimposed images of both stains. **(B)** At 4 days post-infection, cells were sorted into mCherry+GFP- and mCherry+GFP+ populations and quantitative real-time PCR analysis for true late genes UL44 and pp28 was performed. Error bars represent SD of triplicates, statistical analysis performed with Student's *t*-test. *p < 0.05, **p < 0.01, ***p < 0.001, ****p < 0.0001.

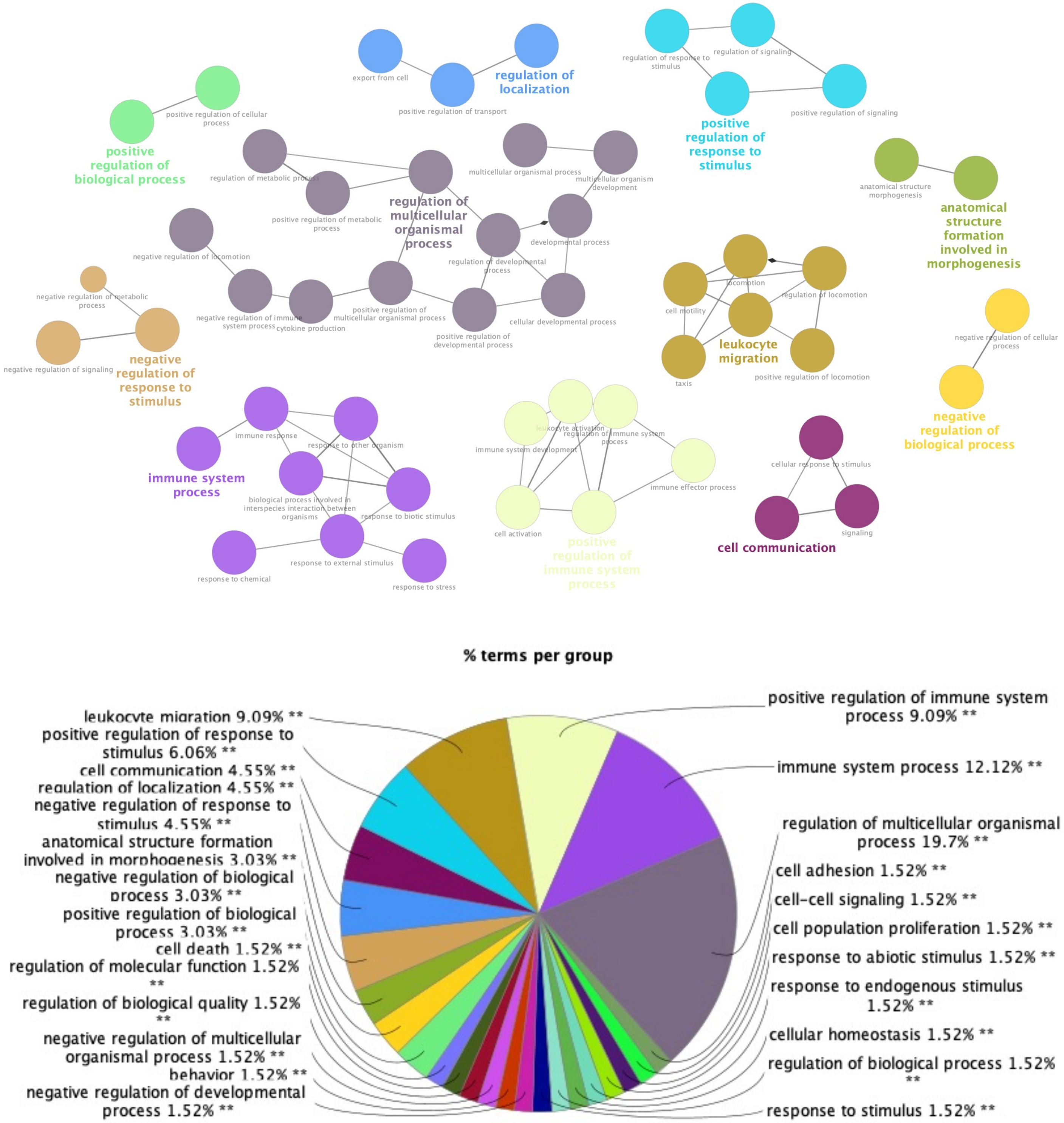


**Supp Fig 2. Gene Ontology analysis of cytokine array performed on secretome from CD4+ cells co-cultured with HCMV-infected fibroblasts compared with secretome from CD4+ cells co-cultured with uninfected fibroblasts**

Gene ontology (GO) analysis using ClueGO$^{TM}$ was performed on the results of the cytokine array performed in Fig 3. to find the GO terms most commonly associated with the cytokines that had been upregulated in each comparison in the assay, after elimination of cytokines that had a p value of >0.05 and a $\log_2$(fold change) value of <1. **(A)** shows the GO terms in bubble plot format, where size of circle corresponds to p value of the association, **(B)** shows the percentages of terms associated with each GO group, and **(C)** shows the amount of cytokines associated with each GO term, expressed as a percentage of total number of cytokines analysed.

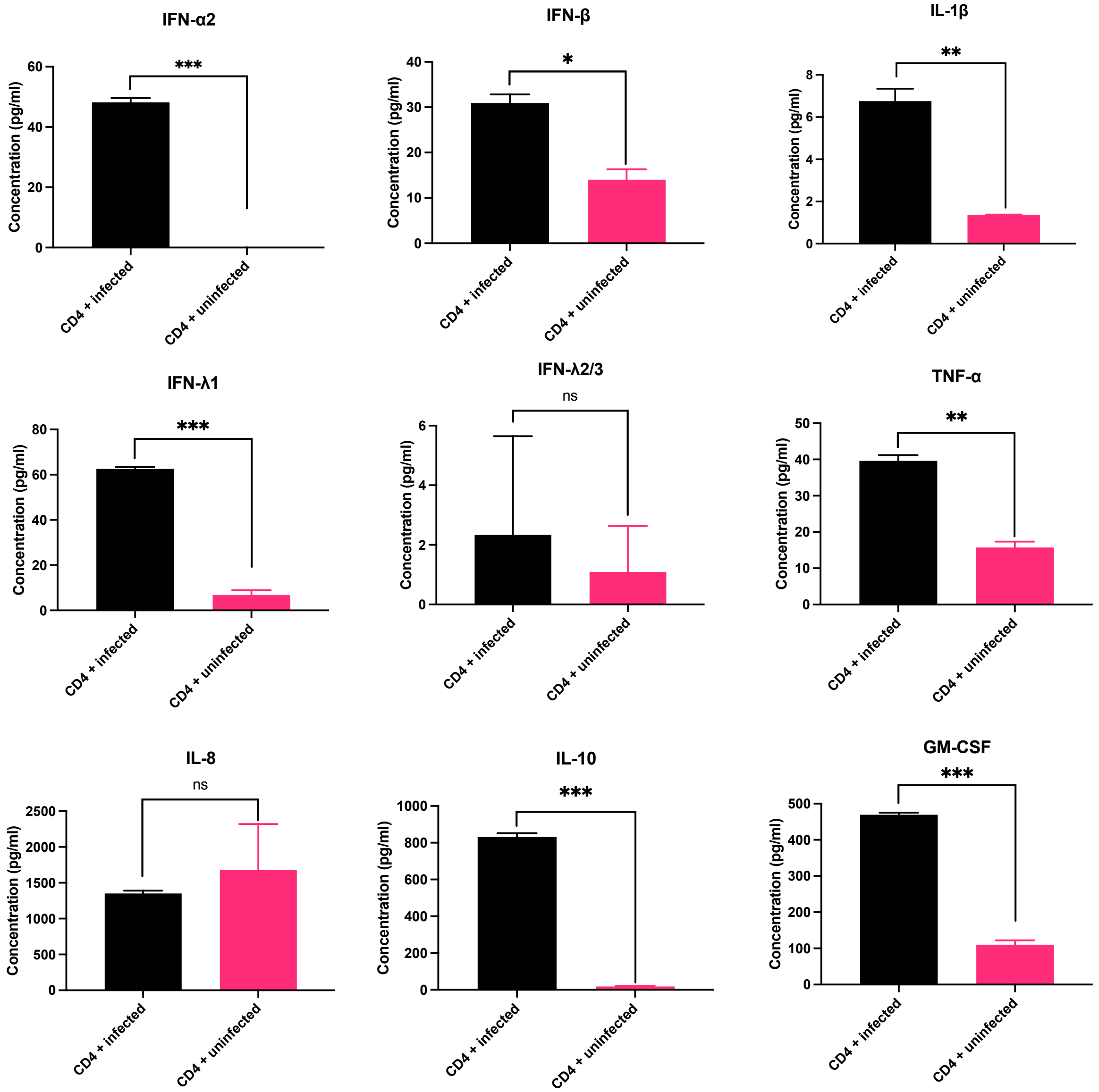


**Supp Figure 3. Concentrations of various cytokines in secretome of CD4+ cells co-cultured with HCMV-infected autologous fibroblasts**

LegendPlex analysis of 9 cytokines was performed on secretomes from CD4+ cells co-cultured with infected autologous fibroblasts (“CD4 + infected”) and uninfected autologous fibroblasts (“CD4 + uninfected”).

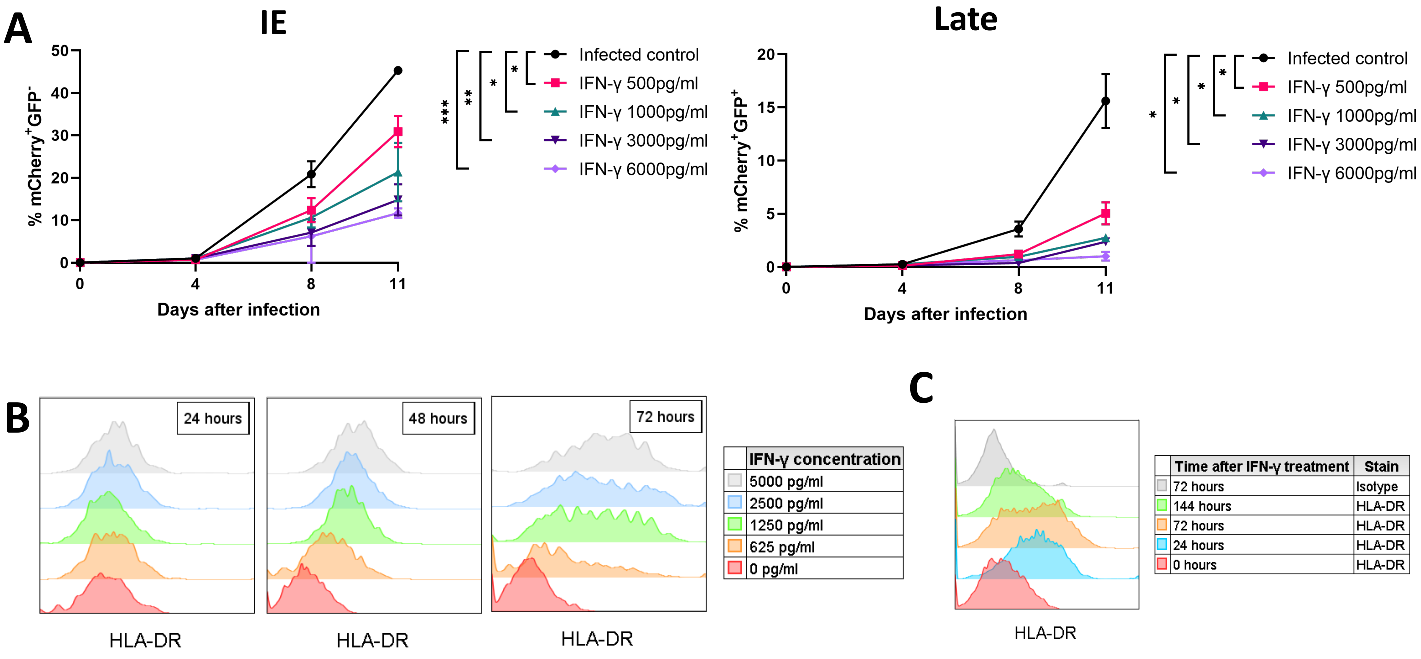


**Supp Figure 4. Effects of IFN-γ on HCMV-infected and uninfected fibroblasts**
**(A)** Recombinant IFN-γ was added to HCMV-infected fibroblasts (MOI=0.03) at 1 dpi at various concentrations and further incubated for 7 days before harvest and analysis by flow cytometry for percentage of cells in IE and late CMV gene expression. **(B)** Recombinant IFN-γ was added to HFFFs at various concentrations shown and incubated for 24, 48 or 72 hours before staining for HLA-DR and analysis by flow cytometry. **(C)** Recombinant human IFN-γ (1000 pg/ml) was added to HFFFs and incubated for durations shown before staining for HLA-DR and analysis by flow cytometry. Error bars represent SD of triplicates, statistical analysis performed with Student's *t*-test. *$p < 0.05$, **$p < 0.01$, ***$p < 0.001$.

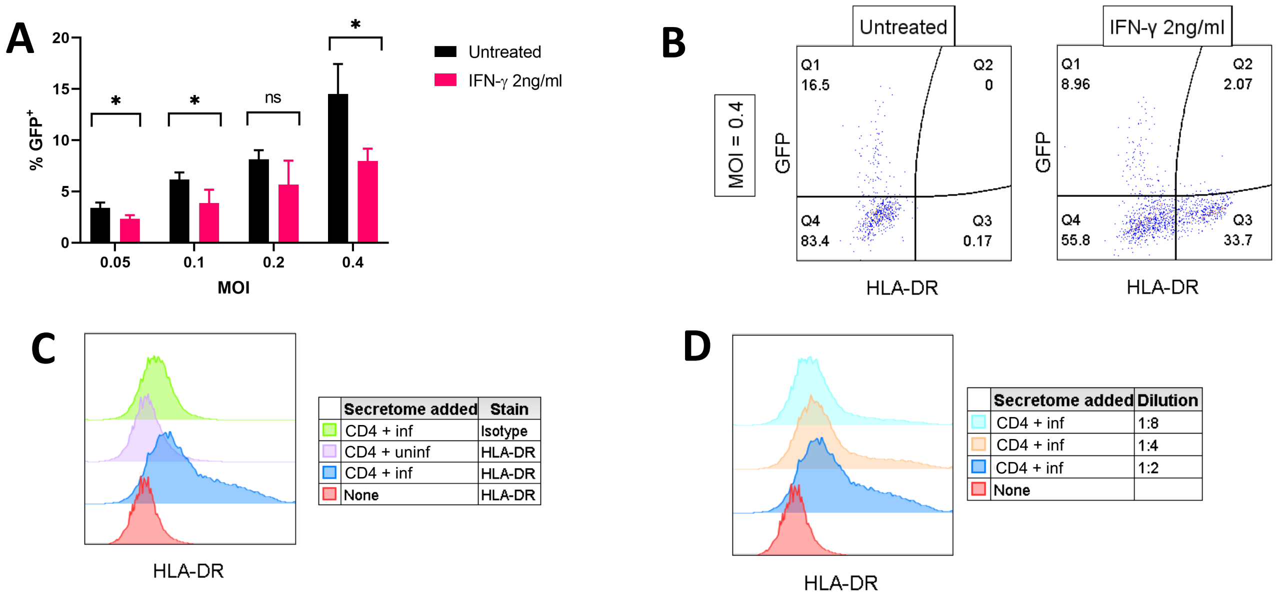


**Supp Figure 5. IFN-γ inhibits late viral gene expression in HUVECs, and secretome of CD4+ cells co-cultured with HCMV-infected fibroblasts induces MHC Class II via on HUVECs and ARPEs**

**(A)** Recombinant IFN-γ (2000 pg/ml) was added to HCMV-infected HUVECs at 1 dpi at MOIs shown. At 6 dpi, IFN-γ was topped up by removing media in the wells and adding fresh media with IFN-γ (2000 pg/ml) or media was changed in the untreated controls. At 12 dpi, wells were harvested and analysed by flow cytometry. Error bars represent SD of triplicate wells, statistics performed using Student's *t*-test. **(B)** Sample of flow cytometry plots from MOI=0.4 of experiment in (A). **(C)** "CD4 + WT" or "CD4 + uninf" secretome was added to adult retinal pigmented epithelium (ARPE-19) cells and incubated for 48 hours before staining for HLA-DR and analysis by flow cytometry. **(D)** "CD4 + WT" secretome was added to ARPE-19 at dilutions shown and incubated for 48 hours before staining for HLA-DR and analysis by flow cytometry. Error bars represent SD of triplicate wells, statistical analysis performed with Student's *t*-test. *$p < 0.05$.
